# A transcription regulatory network within the ACE2 locus may promote a pro-viral environment for SARS-CoV-2 by modulating expression of host factors

**DOI:** 10.1101/2020.04.14.042002

**Authors:** Tayaza Fadason, Sreemol Gokuladhas, Evgeniia Golovina, Daniel Ho, Sophie Farrow, Denis Nyaga, Hong Pan, Neerja Karnani, Conroy Wong, Antony Cooper, William Schierding, Justin M. O’Sullivan

## Abstract

**Introduction:** A novel severe acute respiratory syndrome coronavirus 2 (SARS-CoV-2) was recently identified as the pathogen responsible for the COVID-19 outbreak. SARS-CoV-2 triggers severe pneumonia, which leads to acute respiratory distress syndrome and death in severe cases. As reported, SARS-CoV-2 is 80% genetically identical to the 2003 SARS-CoV virus. Angiotensin-converting enzyme 2 (ACE2) has been identified as the main receptor for entry of both SARS-CoV and SARS-CoV-2 into human cells. ACE2 is normally expressed in cardiovascular and lung type II alveolar epithelial cells, where it positively modulates the RAS system that regulates blood flow, pressure, and fluid homeostasis. Thus, virus-induced reduction of ACE2 gene expression is considered to make a significant contribution to severe acute respiratory failure. Chromatin remodeling plays a significant role in the regulation of ACE2 gene expression and the activity of regulatory elements within the genome.

**Methods:** Here, we integrated data on physical chromatin interactions within the genome organization (captured by Hi-C) with tissue-specific gene expression data to identify spatial expression quantitative trait loci (eQTLs) and thus regulatory elements located within the ACE2 gene.

**Results:** We identified regulatory elements within *ACE2* that control the expression of PIR, CA5B, and VPS13C in the lung. The gene products of these genes are involved in inflammatory responses, *de novo* pyrimidine and polyamine synthesis, and the endoplasmic reticulum, respectively.

**Conclusion:** Our study, although limited by the fact that the identification of the regulatory interactions is putative until proven by targeted experiments, supports the hypothesis that viral silencing of *ACE2* alters the activity of gene regulatory regions and promotes an intra-cellular environment suitable for viral replication.

## Introduction

Within months of the first reports [1], the COVID-19 outbreak has become a pandemic infecting and killing thousands of people worldwide [2]. COVID-19 is an infectious disease associated with acute respiratory distress syndrome (ARDS) that is caused by SARS-CoV-2, a Betacoronavirus that is 80% identical to the SARS-CoV virus [3]. Betacoronaviruses, including SARS-CoV, Murine Hepatic Virus (MHV), and SARS-CoV-2, utilize the ACE2 protein for cell entry [4,5]. The Spike protein on SARS-CoV-2 has a 10 to 20 fold higher affinity for the ACE2 protein than its SARS-CoV homologue [3,6].

The ACE2 protein is highly expressed in cardiovascular and lung type II alveolar epithelial cells [3,7,8], where ACE2 is a primary modulator of the renin–angiotensin (RAS) system that regulates blood flow, pressure and fluid homeostasis [9]. The ACE2 protein and the products of the reactions it catalyzes have also been implicated in immune responses and antiinflammatory pathways [10–12].

SARS-CoV infection reduces *ACE2* gene expression and this is thought to contribute to severe acute respiratory failure [4] by triggering an imbalance in the RAS system that causes a loss of fluid homeostasis, induces inflammatory responses [10,13,14], and results in severe acute injury in heart and lung [3,15,16]. As mentioned above, both SARS-CoV and SARS-CoV-2 utilize the ACE2 protein for cell entry. Poor prognoses in elderly SARS-CoV-2 patients (≥65 years old) are frequently associated with a pre-existing reduction in *ACE2* expression and imbalance in ACE2-related host derived pathways [17,18]. *ACE2* is an X-linked gene whose expression is regulated by chromatin structure. Brg1, a chromatin remodeler, and the FoxM1 transcription factor recognize the *ACE2* promoter and reduce expression through a mechanism involving structural chromatin changes [19]. This control is complex, as illustrated by the finding that *ACE2* gene escapes X chromosome-inactivation and shows a heterogeneous sex bias that is tissue dependent [20].

Chromatin structure in the nucleus involves non-random folding of DNA on different scales [21]. This folding and the resulting contacts that form are dynamic, and can be disrupted (*e.g.* by genetic variation) leading to altered enhancer-promoter interactions that result in changes in gene expression [22]. Changing chromatin structure rewires interactions between regulatory elements and the genes they control. Theoretically and practically, each component of this change contributes to the observed pathogenesis [23,24], and can lead to developmental disorders[25] and cancer [26–28].

Virus-induced chromatin changes at the ACE2 locus could induce expression changes in additional genes regulated by elements located within this locus and thus may alter/modulate host factors important for SARS-CoV-2 replication. How can you identify the elements within a gene regulatory network like this? One approach to identify the networks that form between regulatory elements and the genes they control is to use information on the physical interactions that are captured occurring between the elements. Physical interactions between two sites can be captured and identified using Hi-C [29,30]. We have used this insight to develop a discovery-based pipeline (CoDeS3D; S1 Fig) [23]. Our approach uses genetic variation (*e.g.* single nucleotide polymorphisms) to identify changes in gene expression and thus determine if a region that physically contacts a gene contains a regulatory element. This enables the rapid identification of the regulatory networks that form in cells and tissues (*e.g.* [23,31]).

We hypothesized that *ACE2* and its flanking region contained regulatory elements that coordinate the expression of other genes, and that virus induced chromatin changes at ACE2 inadvertently modulate host factors that promote viral replication. Here we undertook an indepth characterization of the regulatory control regions within *ACE2* and their activity in lung tissue. Regulatory elements located within *ACE2* affect the expression levels of the *PIR* and *CA5B* genes. PIR and CA5B are involved in NF-κB regulation and pyrimidine synthesis, respectively. *VPS13C*, encoding a factor required for late stage endosome maturation, is also controlled by a putative enhancer located in intron 11 of *BMX,* adjacent to *ACE2.* We propose that *ACE2* repression by SARS-CoV-2 trips a chromatin-based switch that coordinates the activity of these regulatory elements and thus the genes they control. Collectively, these changes inadvertently lead to the development of a pro-viral replication environment.

## Methods

### Identification of SNPs in the *ACE* locus

We selected all common single nucleotide polymorphisms (SNPs) from dbSNP (build153) with a minor allele frequency (MAF) > 1% that were located within chrX:15,519,996-15,643,106, which included the *ACE2* gene and its flanking region (hereafter *ACE2* locus). SNP positions are as reported for the human genome build hg38 release 75 (GRCh38).

### Identification of tissue-specific SNP-gene spatial relationships in the *ACE* locus

We used the CoDeS3D algorithm [23] to identify putative spatial regulatory interactions for all SNPs at the *ACE2* locus (S1 Fig). CoDeS3D integrates data on physical chromatin interactions within the genome organization (captured by Hi-C) with tissue-specific gene expression data to identify spatial expression quantitative trait loci (eQTLs). To get lung-specific spatial connections, we identified SNP-gene pairs across lung-specific Hi-C libraries using published data for IMR90, A549, and NCI-H460 cell lines and lung tissue (GEO accession numbers GSE35156, GSE43070, GSE63525, GSE105600, GSE105725, GSE92819, GSE87112, S1 Table). We then queried GTEx for eQTL associations with lung tissue (dbGaP Accession phs000424.v8.p2, https://gtexportal.org/home/). The age of the GTEx lung sample donors peaks between 50-60 years (S2 Fig). SNPs were assigned to the appropriate Hi-C restriction fragments by digital digestion of the hg38 reference genome (matching the restriction enzyme from the Hi-C libraries: *MboI* or *HindIII).* All SNP-fragments were queried against the Hi-C databases to identify the distal DNA fragments with which they physically interact. For each distal fragment, which overlapped a gene coding region, a SNP-gene spatial connection was confirmed. There was no binning or padding around restriction fragments to obtain gene overlap. Spatial tissue-specific SNP-gene pairs with significant eQTLs (both cis-acting [<1Mb between the SNP and gene] and trans-acting eQTLs [>1Mb between the SNP and gene or on different chromosomes]; FDR adjusted p < 0.05) within the lung were subsequently identified by querying GTEx v8 lung tissue (UBERON:0008952).

### URLs

GEO database: https://www.ncbi.nlm.nih.gov/geo/

CoDeS3D pipeline: https://github.com/Genome3d/codes3d-v2

GTEx Portal: https://gtexportal.org/home/

GUSTO study: http://www.gusto.sg/

### Data and code availability

All python and R scripts used for data analysis and visualization are available at https://github.com/Genome3d/ACE2-regulatory-network. R version 3.5.2 and RStudio version 1.2.5033 were used for all R scripts. All python scripts used Python 3.7.6.

## Results

### The *ACE2* locus harbors regulatory variants that control SARS-CoV-2 relevant cellular functions

We tested 367 common SNPs located across the *ACE2* locus (chrX: 15,519,996-15,643,106) for their potential to act as spatial eQTLs. None of the common SNPs we tested affected *ACE2* expression levels in lung tissue (S2 Table).

The wider *ACE2* locus (chrX: 15,519,996-15,643,106; GRCh38/hg38) sits within a topologically associating domain (TAD) that is conserved across some tissues, *e.g.* IMR90 (Fig 1A). Therefore, it was not surprising that we identified control elements within this *ACE2* locus (Fig 1A). The distribution of targets for the putative control elements we identified is consistent with previous studies that show that while the majority of significant eQTLs fall within 100 kb of the transcription start site of a gene, only 60% of all eQTLs are upstream of the gene they regulate[32]. Notable amongst the elements we identified are long distance *trans-regulatory* interactions involving: 1) rs1399200:*VPS13C* (chr15:61,852,389-62,060,473; encodes vacuolar protein sorting-associated protein 13C); and 2) *rs6632680:PHKA2* (chrX:18,892,300-18,984,598; encodes phosphorylase kinase regulatory subunit alpha 2) (S2 Table).

**Fig 1.**
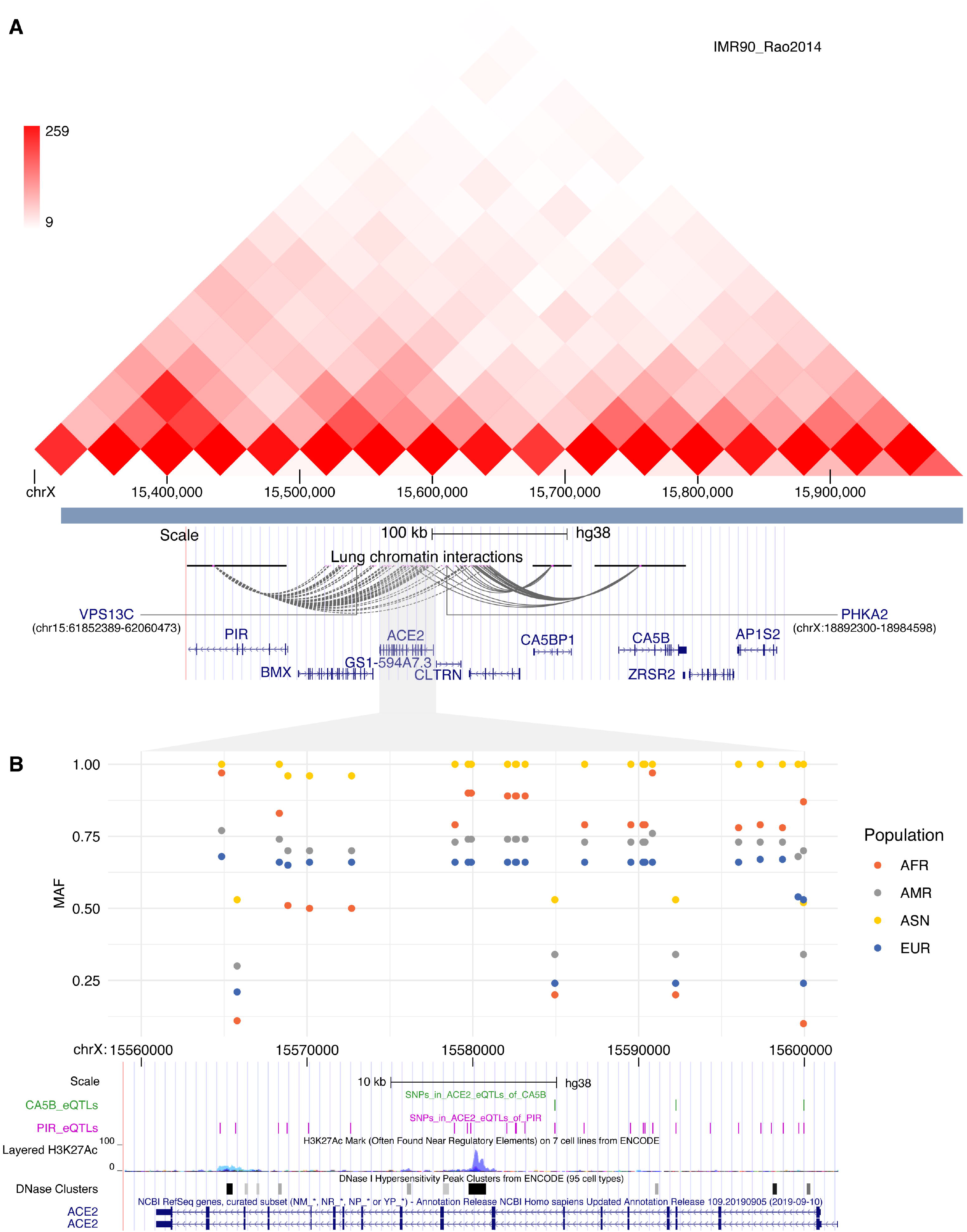
Elements located within and surrounding the *ACE2* locus regulate the lung-specific expression of *PIR, CA5B, CA5BP1, VPS13C,* and *PHKA2.* (A) Common genetic variants (SNPs) located within the *ACE2* locus form spatial *cis*-acting regulatory interactions with *PIR*, *CA5BP1,* and *CA5B* across sub-TAD boundaries on chrX:15,300,000-15,600,000. Inter-TAD *trans-acting*interactions regulate *PHKA2* (3.2 Mb away) and *VPS13C*(located on chromosome 15). Visualization of TAD and chromatin interactions was performed using the 3D genome browser (http://yuelab.org/)[35] and UCSC browser’s interact tool (http://genome.ucsc.edu)[36], respectively. (B) Within *ACE2,* MAFs for the SNPs that tag the regulatory sites showed significant bias in four different populations (*i.e.* African [AFR], Ad Mixed American [AMR], East Asian [ASN] and European [EUR]) at one *PIR* (rs714205) and three *CA5B* regulatory sites (rs4646142, rs2285666, and rs2106809), consistent with selection acting on these loci. MAFs were obtained from HaploReg v4.1 (https://pubs.broadinstitute.org/mammals/haploreg/haploreg.php) [37].

We identified eighty genetic variants within the ACE2 locus as *cis*-acting spatial eQTLs that physically modulate the expression of genes *PIR* (encodes Pirin), *CA5BP1* (a pseudogene of CA5B), and *CA5B* (encodes mitochondrial carbonic anhydrase) in lung tissues (S2 Table). Fifty-eight SNPs located across the region are associated with increased expression of *PIR* (log2[aFC, allelic fold change] 0.462 ± 0.07) consistent with the elements they mark repressing PIR transcription. Eighteen SNPs located across the region are associated with decreased expression of *CA5B* (log2[aFC] −0.257 ± 0.005) consistent with the elements they mark enhancing *CA5B* transcription. These variants occurred in two clusters: 1) within the *ACE2* gene; and 2) within the *CLTRN*(*TMEM27*) gene – a known homologue of *ACE2.* Expression of *CA5BP1,* a pseudogene of *CA5B,* was also repressed (log2[aFC] −0.21 ± 0.01) by 6 SNPs within the *ACE2* locus. Within *ACE2* itself there were only control regions for the *PIR* and *CA5B* genes (Fig 1B).

The common variants that we tested show an unusual ancestry associated pattern of minor allele frequencies (Fig 1B). Specifically, the East Asian population (1K Genomes project) displays little variation across the bulk of the variants we analyzed. This observation is supported by measures of genetic diversity (FST) between the Indian, Chinese and Malay populations within the Growing Up in Singapore Towards healthy Outcomes (GUSTO) cohort (S3 Table). However, this pattern breaks down at several positions across the *ACE2* gene (including rs4646142, rs2285666, and rs2106809, which show significant selection towards the reference allele) in all of the tested populations, indicating potential selective pressure at these loci (Fig 1B). Notably, two of these variants alter potential transcription factor binding sites (*i.e.* rs2285666 alters HNF1, and Ncx motifs, rs2106809 alters a CEBPB motif; S4 Table). All three variants (rs4646142, rs2285666, and rs2106809) have previously been associated with allele, sex and ethnicity specific impacts on hypertension, blood pressure, hypertrophic cardiomyopathy, type 2 diabetes, myocardial infarction (reviewed in[33]). Moreover, the CEBPB motif is recognized by the CCAAT enhancer binding protein-β which has been implicated in inflammatory responses in lung carcinoma cells [34].

## Discussion

We identified transcription regulatory elements for *CA5B* and *PIR* that are active in lung tissue and are located within the *ACE2* gene. We also identified a transcription regulatory element (located in the *BMX* gene, adjacent to *ACE2*) for the *PIR* and *VPS13C* genes. It is sterically impossible for a single DNA sequence to simultaneously be transcribed and regulate another gene through a physical connection. Therefore, we propose that SARS-CoV-2-induced chromatin-dependent repression of *ACE2* expression in lung enables the regulatory sites, repressing *PIR* and activating *CA5B,* to exhibit increased functionality in infected cells (Fig 2). We hypothesize that this regulatory change extends to coordinate changes in the expression of *VPS13C* and *PHKA2* in ways that promote viral proliferation. This host regulatory network has not evolved to benefit the virus but rather, these regulatory changes inadvertently produce an environment advantageous for the virus.

**Fig 2.**
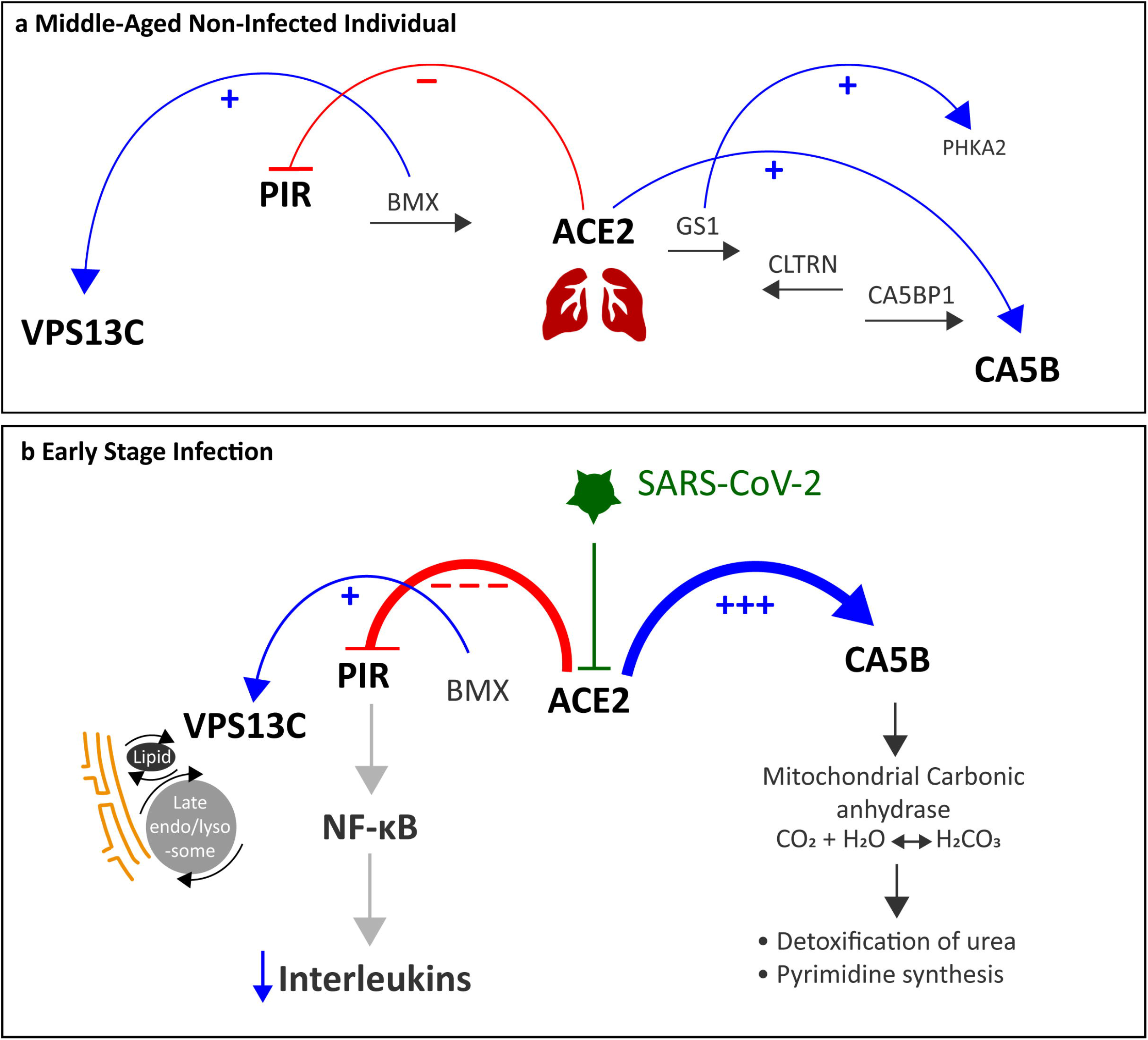
Hypothesis: SARS-CoV-2 infection is associated with an *ACE2* dependent switch that alters expression of proteins that promote an environment for viral proliferation in the lung. (A) In middle-aged non-infected individuals, control regions within *ACE2* are capable of downregulating the expression of *PIR*, which is involved in the NF-κB pathway. Enhancer elements within *ACE2* are poised to upregulate *CA5B* expression, which encodes an enzyme important for pyrimidine synthesis. In addition to this, an enhancer region within the *BMX* gene (still within the same TAD) contributes to *VPS13C* regulation. (B) We hypothesize that upon viral infection, SARS-COV-2 represses *ACE2* expression, which increases the activity of the *PIR* repressor and *CA5B* enhancer. This results in a reduction in the production of PIR – the redox switch necessary for NF-κB activation, while also increasing pyrimidine synthesis, which is necessary for viral replication.

The *CA5B* gene encodes a mitochondrial carbonic anhydrase that catalyzes the reversible hydration of CO_2_ in the lung. This reaction is important in mitochondria as it supplies HCO_3_^-^ ions required by pyruvate carboxylase for gluconeogenesis, and by carbamoyl phosphate synthase 1 (CSP1) for pyrimidine biosynthesis (Fig 2B) [38–40]. Pyrimidines are important host factors critical for viral genomic replication, mRNA synthesis for protein translation, and phospholipid synthesis [41]. Inhibiting *de novo* pyrimidine biosynthesis impacts on SARS-CoV-2 replication [18]. CSP1 additionally produces a precursor for the biosynthesis of polyamines, small aliphatic molecules that play important roles in virus replication. Inhibition of polyamine biosynthesis significantly impaired replication of the Middle East Respiratory Syndrome [MERS] coronavirus [42]. Targeted inhibition of CA5B encoded carbonic anhydrase might therefore decrease levels of critical host factors pyrimidines and polyamines – critical host factors needed for SARS-CoV-2 replication. Intriguingly, similarities in the pathologies of SARS-CoV-2 infection and high altitude pulmonary edema (HAPE) have led to the suggestion that carbonic anhydrase inhibitors could be used to treat or prevent Covid-19 infection [43].

Interleukin expression is responsible for irreversible, pathological changes associated with SARS-CoV infection in the lung (*e.g.* [44]). Human coronavirus has been shown to fine-tune NF-κB signaling [45]. *PIR* encodes a non-heme iron binding protein that is a redox switch that modulates the binding of p65 (RelA) to NF-κB responsive promoters [46]. NF-κB regulates multiple immune function aspects, including the production of pro-inflammatory cytokines[47]. Therefore, it is notable that repressor regulatory sequences for *PIR* sit within the *ACE2* gene (Fig 2A). We postulate that the chromatin modifications that silence *ACE2* expression upon early stage infection activate the *PIR* repressor (Fig 2B). This reduces responsiveness of NF-κB, and thereby delays the expected and needed anti-viral response. Reduction in PIR expression would also reduce the impact of any changes to intra-cellular redox state caused by that SARS-CoV-2 infection, however little is known and future experiments are required to clarify this.

The enveloped Betacoronaviruses (MHV, SARS-CoV, SARS-CoV-2) gain entry to the cell through the endo/lysosomal pathway and require late endosomal maturation for fusion [48]. Therefore, it is interesting to speculate on the impact of coordinated changes to *VPS13C* expression. The VPS13 family are endoplasmic reticulum associated lipid transporters. VPS13C is proposed to act as a lipid transporter at organelle contact sites between (i) the endoplasmic reticulum (ER) and endolysosomes, and (ii) the ER and lipid droplets, where it transfers lipids, potentially bulk lipid transfer, between organelles to maintain lipid homeostasis and organelle functionality [49]. Increased *VPS13C* expression is predicted to increase the extent of contact and lipid transfer between these organelles. This in turn could enhance the virus’s replication capacity and pathogenesis, as the ER plays both a physical and functional central role in the virus’s capacity to replicate and form new viral progeny. Moreover, SARS-CoV extensively reorganizes the host cell’s membranes infrastructure to produce a reticulo-vesicular network of modified ER to coordinate its replication cycle [50]. Alterations to ER-lipid droplet contacts mediated by VPS13C could support the virus’s required expansion and re-organization of ER membranes by altering lipid flow through the ER [51]. Notably, cells infected with Hepatitis C virus (HCV), a positive-strand RNA virus like SARS-CoV-2, contain ER-derived membranous structures that contain significantly high levels of cholesterol, despite the ER of uninfected cells possessing relatively low cholesterol levels [52]. Increased VPS13C-mediated ER-endolysosomal contact sites could increase the capacity of endocytosed dietary cholesterol to be delivered to the ER and enhance the virus’s ability to replicate. Pharmacological impairment of endolysosomal cholesterol efflux reduced HCV replication, [52] suggesting another possible therapeutic approach for investigation to slow SARS-CoV-2 replication. ER stress, impacted by changes to *VPS13C* expression, may also contribute to late infection stage NF-κB activation (reviewed in [53]).

The significance of the putative enhancer for *PHKA2,* which encodes the phosphorylase kinase regulatory subunit alpha 2, is unclear. Mutations in this gene have been linked to glycogen storage disorders and glucose metabolism. Thus, linkages can be drawn to the increased expression of *CA5B,* which impacts on gluconeogenesis. Notably, *PHKA2* was downregulated in plasma from individuals with hepatocellular-carcinoma caused by HCV infections [54]. Theoretically, chromatin remodeling in response to SARS-COV-2 infection could down-regulate *PHKA2* expression. However, there is a paucity of information linking this gene to viral infections or the lung and this conclusion requires additional experimental support.

SARS-CoV is known to repress *ACE2* expression [4]. *ACE2* regulation involves chromatin remodeling and structural chromatin changes [19]. Several of the regions that we identified overlapped or were adjacent to CTCF biding sites (*e.g.* rs1399200, which regulates *PIR* and *VPS13C;* rs6629111, which regulates *CA5B;* and sites [rs714205, rs1514280, rs4240157 and rs4646131] within *ACE2*[33]). We also note that the regulatory sites we identified included transcription factor binding sites for Ap-1, RXRA (a DNA-binding receptor involved in host-virus interactions) [55], GR or NR3C1 (a regulator of inflammation in asthma and COPD) [56], Pou2f2 (trans-activator of NR3C1)[57], and P300 (a chromatin modifier; S4 Table) [58]. Expression data within the search-based exploration of expression compendium (SEEK) supports a strong co-expression relationship between *ACE2* and *PIR* (lung cancer, ovarian tumor) and a weaker association with *CA5B* (ovarian cancer) (http://seek.princeton.edu/ [59]). However, the possible mechanism(s) that link ACE2 silencing to alterations associated with these regulatory regions remains unknown until empirically determined in lung cells in the presence/absence of real or simulated viral infection.

Whilst this study is novel and uses empirically derived data in the analyses, the observations are limited by the fact that the identification of the regulatory interactions is only putative until proven by targeted experiments. Ideally, the Hi-C datasets should be derived from matched tissues prior and post SARS-CoV-2 infection. Finally, the GTEx database has recognized limitations, including the ethnic diversity of the samples. These limitations will form the basis of future studies.

## Conclusion

We identified putative regulatory regions in and surrounding *ACE2* that regulate the expression of *PIR*, *CA5B,* and *VPS13C* in the lung. We contend that viral induced chromatin-dependent repression of the *ACE2* gene increases the activity of these regulatory sites and promotes an intra-cellular environment suitable for viral replication. The altered gene products represent new targets for anti-SARS-CoV-2 therapeutics.

## Acknowledgements

The authors would like to thank the Genomics and Systems Biology Group (Liggins Institute), Mark Hampton, and Elizabeth Ledgerwood for useful discussions. We would like to thank the funders of GTEx Project – common Fund of the Office of the Director of the National Institutes of Health, and by National Cancer Institute, National Human Genome Research Institute, National Heart, Lung, and Blood Institute, National Institute on Drug Abuse, National Institute of Mental Health, National Institute of Neurological Disorders and Stroke. The authors thank the GUSTO study group for comments on the manuscript. This work has been released as a pre-print.

## Author contributions

TF, EG, SG and WS performed analyses and co-wrote the manuscript. DH, DN, SF, and AC performed literature searches and co-wrote the manuscript. HP and NK provide FST data from GUSTO and commented on the manuscript. CW reviewed the findings and commented on the manuscript. JOS led the study and co-wrote the manuscript.

## Supplementary Tables

**S1 Table.** Lung-specific Hi-C libraries used in the analysis

**S2 Table.** Lung-specific spatial SNP-gene relationships in the *ACE* locus

**S3 Table.** Genetic diversity estimate (Fst) across *ACE2* in the Indian, Malay and Chinese populations in the GUSTO cohort.

**S4 Table.** The common variants overlap DNA binding motifs. Data from Haploreg v4.1 (3/3/2020)

## Supplementary Figures

**S1 Figure. The CoDeS3D algorithm used in this study.** Restriction fragments containing SNPs located within the *ACE* locus (chrX:15,519,996-15,643,106) were identified. Lungspecific Hi-C libraries were interrogated to identify genes in fragments that spatially interact (in cis- and trans-) with SNP-containing fragments. The identified spatial SNP-gene pairs were further used to query GTEx lung tissue (dbGaP Accession phs000424.v8.p2, UBERON:0008952). The Benjamini-Hochberg FDR control algorithm was applied to adjust the p values of the resulting eQTL associations to identify only significant (FDR < 0.05) lung-specific SNP-gene spatial relationships in the *ACE* locus.

**S2 Figure. The eQTL data used in this study was obtained from lung samples taken from middle-aged individuals.** To assess the correlation of genetic variation with the changes in gene expression, the GTEx project (https://gtexportal.org/home/) collected and analysed lung samples from donors who were densely genotyped. The age-distribution graph illustrates that approximately 70% of the lung samples that were obtained were from donors aged between 50 and 60.

