## Supplementary Figures for "A transcription regulatory network within the ACE2 locus may promote a pro-viral environment for SARS-CoV-2 by modulating expression of host factors"

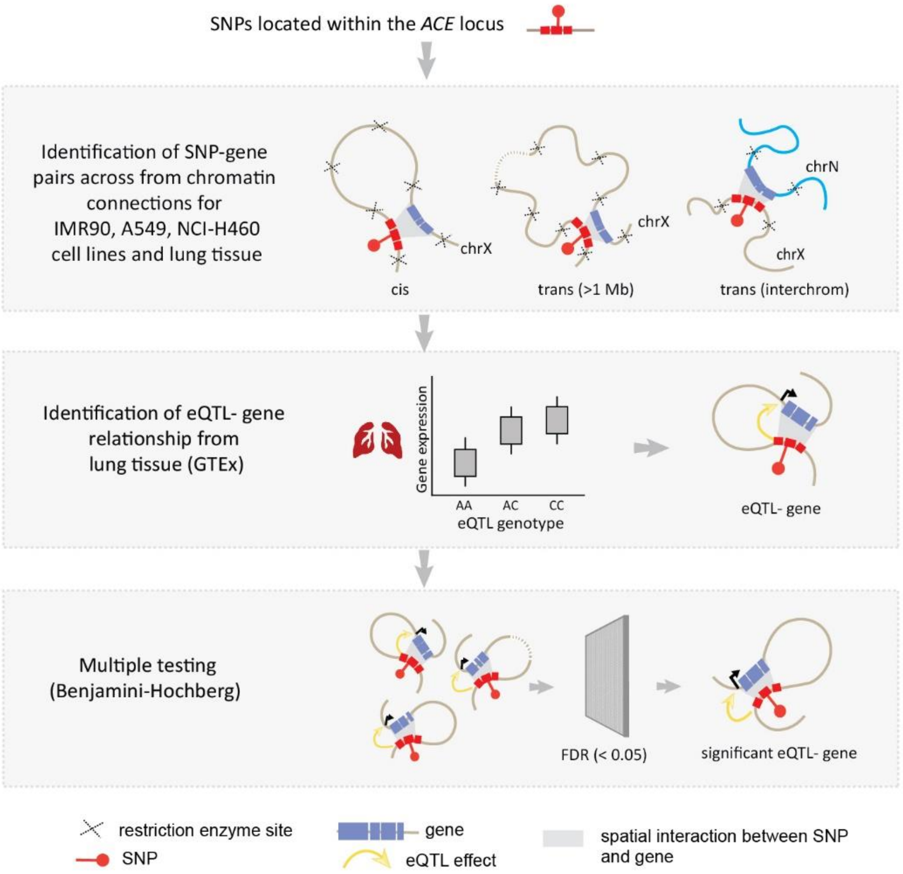


S1 Figure. **The CoDeS3D algorithm used in this study**. Modified from Gokuladhas *et al.* 2020 [1].

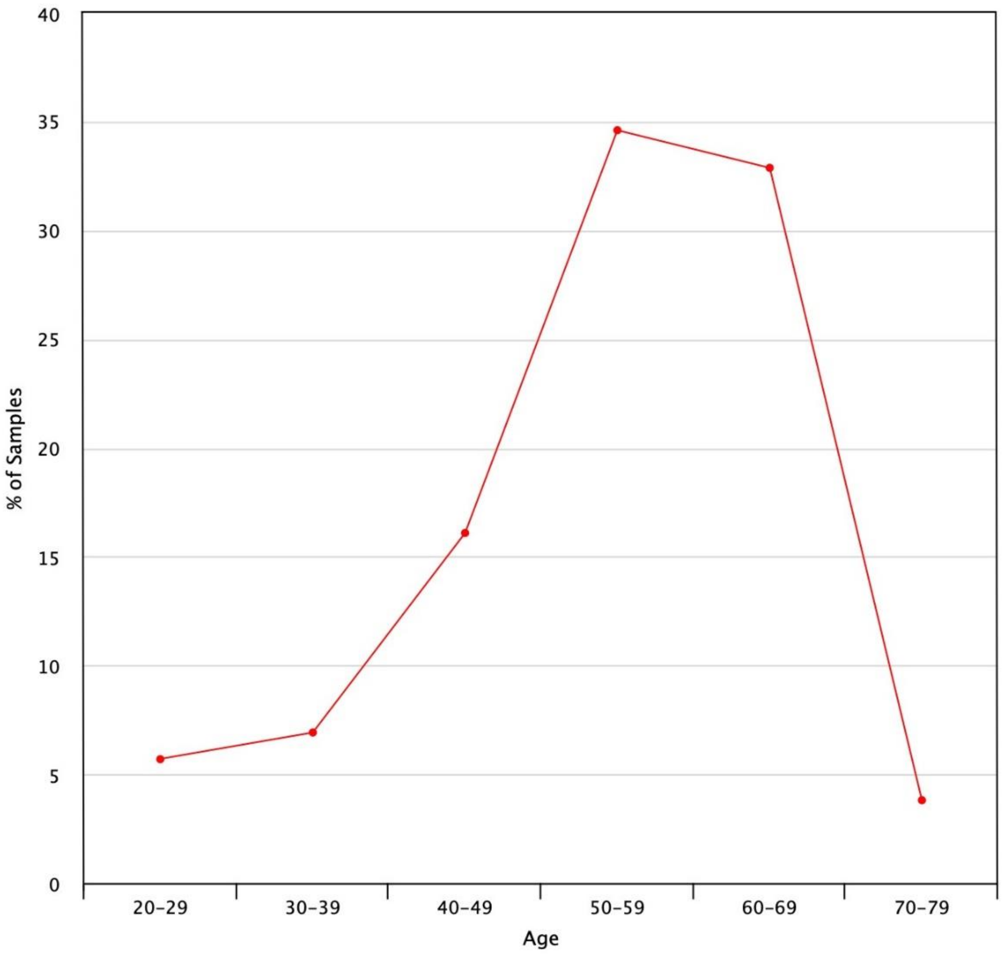


**S2 Figure. The eQTL data used in this study was obtained from lung samples taken from middle-aged individuals.** To assess the correlation of genetic variation with the changes in gene expression, the GTEx project (https://gtexportal.org/home/) collected and analysed lung samples from donors who were densely genotyped. The age-distribution graph illustrates that approximately 70% of the lung samples that were obtained were from donors aged between 50 and 60.
